# Novel Function of Arylalkalamine N-acetyltransfe- rase-1 in Modulating Host Coagulation and Blood Feeding in *Aedes aegypti*

**DOI:** 10.1101/2025.08.27.672538

**Authors:** Xue Gong, Dingfeng Duan, Xiaojing Zhu, Yaneng Huang, Jiukai Chen, Linlong Jiang, Lei Zhang, Qian Han, Chenghong Liao

## Abstract

During blood feeding, mosquitoes introduce substantial quantities of salivary proteins into their hosts to facilitate feeding. Arylalkylamine N-acetyltransferase-1 (aaNAT1) is a salivary protein prominently produced in the salivary glands of *Aedes aegypti*. Previous research has demonstrated that mosquitoes with *AaaaNAT1* mutations fail to efficiently inhibit host blood coagulation; however, the precise regulatory mechanisms remain unknown. This study identified the primary metabolites and proteins interacting with AaaaNAT1 through integrated transcriptomic, proteomic, and metabolomic analyses. Combined with isothermal titration calorimetry (ITC), immunohistochemistry, ELISA, platelet function assays, and coagulation tests, we further established that *Aa*aaNAT1 binds norepinephrine, reducing host platelet aggregation and thereby limiting activation of the host coagulation cascade and fibrinolytic system. Additionally, behavioral investigations showed that *AaaaNAT1* knockdown in *Aedes aegypti* significantly prolonged the time required for female mosquitoes to locate a host and successfully obtain blood, while also markedly reducing blood consumption. This effect may stem from the inability of *aaNAT1*-depleted mosquitoes to effectively suppress host blood aggregation during feeding, as well as the downregulation of octopamine following *AaaaNAT1* knockdown. Therefore, this study reveals a novel role for *aaNAT1* distinct from its known functions in pigmentation and immunity, advancing our understanding of *aaNAT1*’s role in insects.

## Introduction

*Aedes aegypti*, a species in the genus Aedes (family Diptera, phylum Arthropoda), is widely recognized as one of the most dangerous mosquito species due to its broad presence in tropical and subtropical regions and its role as the primary vector for dengue, Zika, and yellow fever viruses[1]. The reproductive cycle of female *A. aegypti* requires hematophagy, as blood from humans or animals is essential for ovarian development and reproduction. During blood feeding, a hemostatic tripod-comprising vasoconstriction, platelet aggregation, and blood flow regulation is triggered [2]. To counteract this, female mosquitoes inject anticoagulant salivary proteins into the host to facilitate blood intake [3].

Arylamine acetyltransferases (aaNATs) catalyze the transfer of an acetyl group from acetyl-CoA to arylamines[4]. In animals, aaNATs regulate melatonin synthesis by acetylating serotonin, thereby influencing circadian rhythms[5–7]. They also catalyze the formation of acetyl-dopamine, which plays a key role in immunity[8–10]. Consequently, studies on insect aaNATs, particularly in *Drosophila*, have focused on their roles in melanization and immunity[11,12]. In *Aedes aegypti*, thirteen *aaNAT* genes have been identified[13], each with distinct functions across tissues, using various biogenic amines as substrates [4,13–17]. Among these, *AaaaNAT1* has broad substrate specificity and plays a critical role in mosquito physiology. Previous studies suggest that *AaaaNAT1* helps detoxify by modulating reactive oxygen species (ROS) levels in the midgut[18]. Furthermore, silencing *AaaaNAT1* impairs the anticoagulant function of salivary glands in female mosquitoes[16]. The results suggested a previously unrecognized function of *AaaaNAT1* in anti-coagulation. However, during mosquito feeding, the host activates coagulation mechanisms-including platelet aggregation[19,20], vasoconstriction[21], and blood coagulation[22]-to resist blood loss, which specific phase *AaaaNAT1* affects and how it exerts this control remain unclear and warrant further investigation.

This study generated a knockdown strain of *AaaaNAT1* in *Aedes aegypti* using RNAi technology, leading to the collection of 40,000 pairs of salivary glands from both the control group (treated with *GUS* dsRNA) and the interference group (*AaaaNAT1* knockdown). We conducted comprehensive transcriptomic, metabolomic, and proteomic analyses of the salivary glands to identify key metabolites and proteins associated with *AaaaNAT1* function. Based on the omics results, *in vitro*–prepared AaaaNAT1 protein was used to determine substrate binding preference via isothermal titration calorimetry (ITC). The impact of AaaaNAT1 on host coagulation was subsequently evaluated using four coagulation assays, platelet function tests, immunohistochemistry, and ELISA. AaaaNAT1 was shown to effectively clear norepinephrine from wound tissue, delay platelet aggregation, and inhibit both the coagulation cascade and fibrinolysis following platelet implantation in mice. Behavioral studies revealed significant differences in blood-feeding behavior between *AaaaNAT1*-knockdown mosquitoes and the control group. This study combined *in vitro* and *in vivo* experiments with molecular and behavioral analyses to preliminarily uncover the role of AaaaNAT1 in enhancing mosquito anticoagulant activity and modulating blood-feeding behavior. By elucidating this previously unknown anticoagulant mechanism, the findings improve our understanding of mosquito–host interactions and may aid in identifying novel intervention targets and strategies, ultimately strengthening efforts to control mosquito-borne viral diseases. Further research into the anticoagulant properties of *AaaaNAT1* may also offer new insights into thrombosis prevention and treatment.

## Results

### 1. RNAi and the Phenotype of Female Mosquito Salivary Glands

Adult mosquitoes that had emerged three days earlier were selected and administered a mixture of bacterial solution containing *AaaaNAT1* dsRNA[23] and sugar water, constituting the interference group. The negative control group consisted of mosquitoes that consumed the same mixture with *GUS* dsRNA, while the blank control group received only sugar water. After eight days of continuous interference, microscopic examination revealed greater atrophy and reduced content in the salivary glands of female mosquitoes in the interference group compared to the two control groups (Fig. 1A). Female mosquitoes from all three groups were collected to assess salivary gland interference efficacy. qPCR results confirmed that *AaaaNAT1* expression was significantly downregulated in both the salivary glands and whole bodies of the interference group (Fig. 1B), indicating effective interference and suitability for omics analysis.

**Fig. 1.**
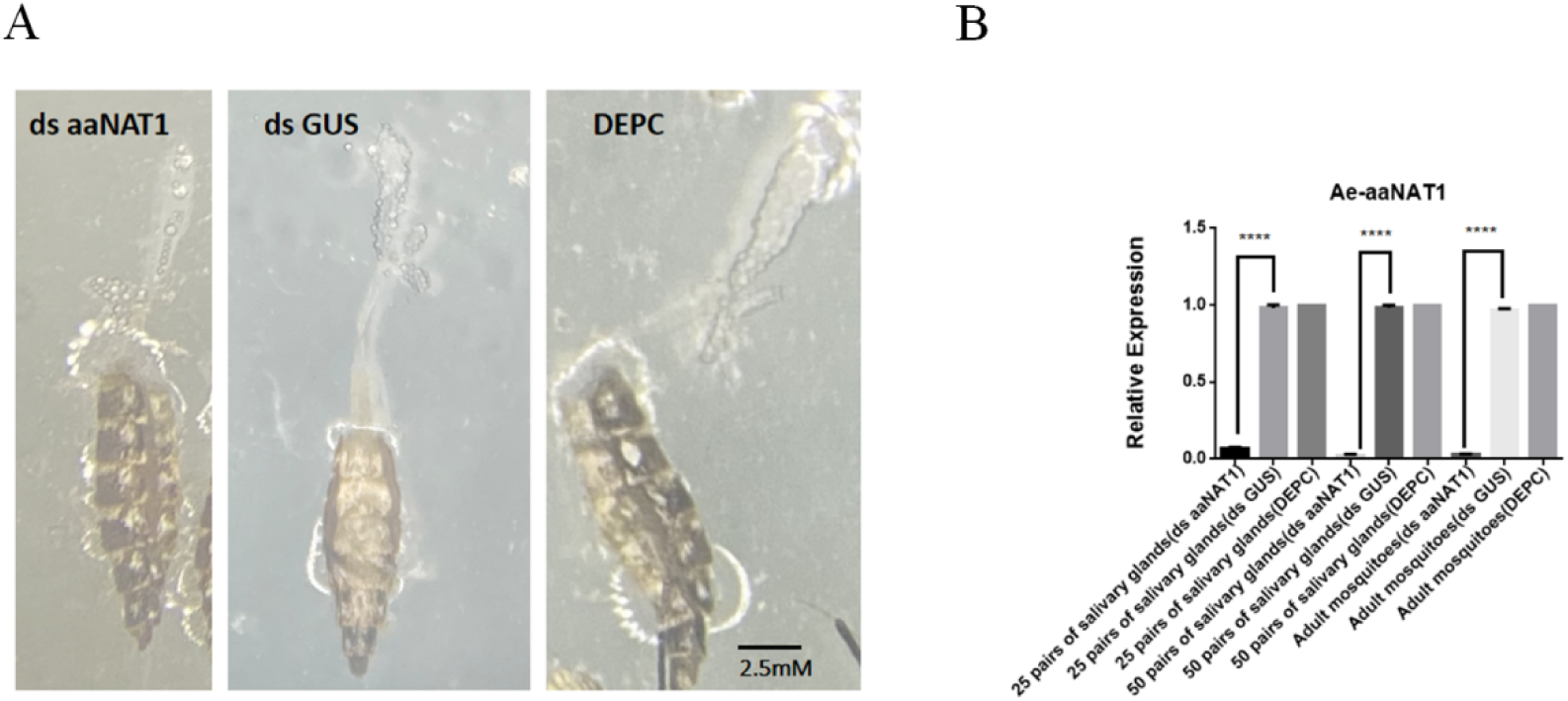
The morphology of mosquito salivary glands after interference and the efficiency of the interference. (**A**) The salivary glands of adult mosquitoes fed with *aaNAT1* dsRNA (ds aaNAT1) appear more shriveled compared to those fed with *GUS* dsRNA (ds GUS) and the blank control (DEPC). (**B**) In adult mosquitoes fed with *aaNAT1* dsRNA (ds aaNAT1), the expression levels of *aaNAT1* in both the 25 and 50 salivary glands, as well as in the whole adult mosquito, were significantly lower than in the groups fed with *GUS* dsRNA (ds GUS) and the blank control group (DEPC). \*\*\*\**p* < 0.0001.

### 2. Significant changes occurred in the three groups after the interference

For transcriptomic, proteomic, and metabolomic analysis, 20,000 pairs of female mosquito salivary glands continuously administered *AaaaNAT1* dsRNA or *GUS* dsRNA for eight days were acquired, with those showing high interference efficiency randomly selected. Several significantly altered genes from the transcriptome were validated by qPCR, with results consistent with transcriptomic data (Fig. S1). Principal component analysis (PCA) revealed a partial separation trend between the *AaaaNAT1* dsRNA and GUS dsRNA groups (Fig. 2A–C). In the metabolome, 2,110 metabolites were identified, with 120 showing significant alterations (Fig. 2D). A total of 6,607 protein/peptide types were identified in the proteome, of which 131 were substantially altered (Fig. 2E). A total of 10,447 genes were identified in the transcriptome, with 52 exhibiting significant changes (Fig. 2F). Altered genes were primarily enriched in the oxidative phosphorylation pathway (Fig. S2), which can induce reactive oxygen species (ROS)[24,25], supporting previous findings that *AaaaNAT1* modulates *Aedes aegypti* detoxification via midgut ROS levels[18]. Furthermore, triomic alterations were concentrated in key pathways linked to *Aedes aegypti* survival, including cofactor biosynthesis, amino sugar and nucleotide sugar metabolism, neuroactive ligand-receptor interactions, pentose and glucuronic acid conversion, and lysosomal and tryptophan metabolism (Fig. 3A). These changes may lead to cellular dysfunction, potentially affecting overall health and survival rates (Fig. 3B).

**Fig. 2.**
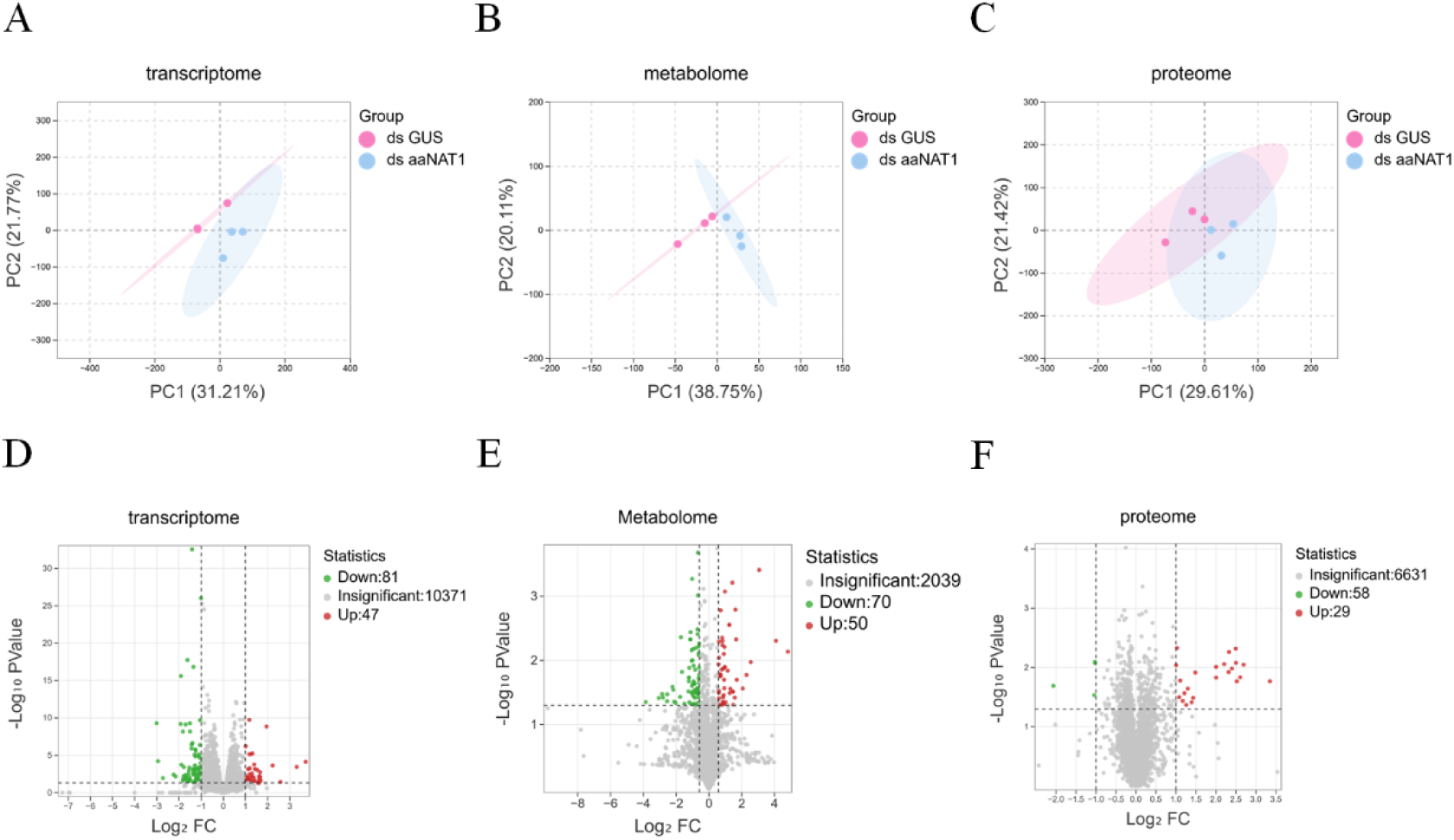
Omics PCA and differential substance volcano plot. (**A**) PCA of the proteome. (**B**) PCA of the metabolome. (**C**) PCA of the transcriptome. (blue) ds aaNAT1: the group fed with *aaNAT1* dsRNA; (red) ds GUS: the group fed with *GUS* dsRNA. (**D**) Volcano plot of the proteome. (**E**) Volcano plot of the metabolome. (**F**) Volcano plot of the transcriptome. Red indicates upregulation, while green indicates downregulation. A deeper color represents a larger fold change.

**Fig. 3.**
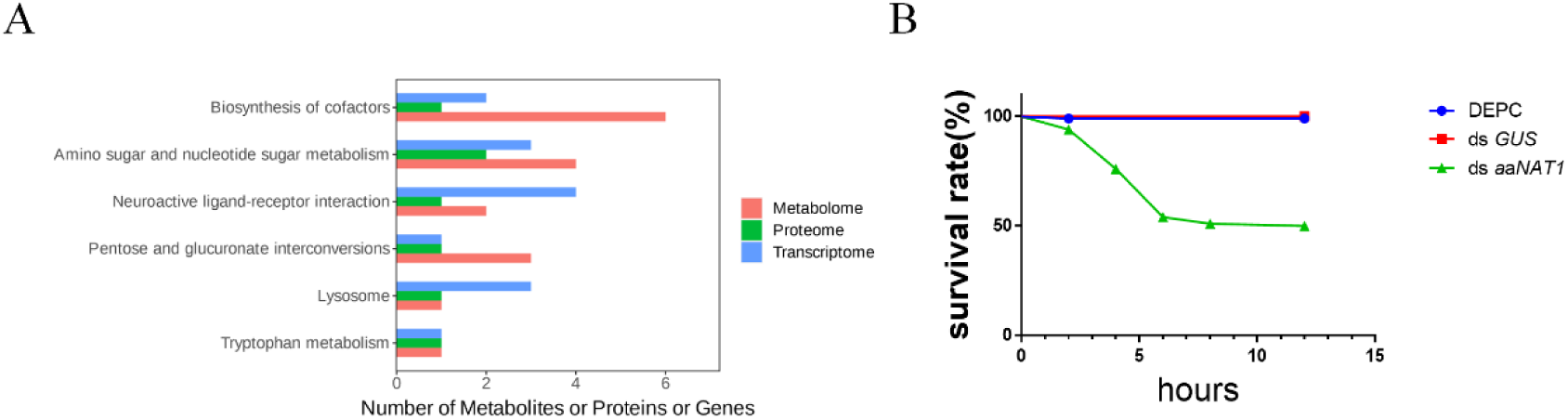
Pathway enrichment map and survival rate curve. (**A**) The six pathways showing the most changes in the transcriptome, proteome, and metabolome. (**B**) Survival curve of *Aedes aegypti* after knockdown of *AaaaNAT1*.

### 3. *AaaaNAT1* affects blood coagulation

The protein compensation effect refers to a phenomenon in which mutations or deletions of specific proteins or genes in an organism prompt other proteins or genes to adaptively change, thereby preserving the organism’s functionality and stability. Upon reviewing the proteomics results, we were surprised to find that among the 72 elevated proteins, some were associated with blood coagulation, including SerpinB5, Serpin27a, AeD7L2, apyrase, and adenosine deaminase (Fig. 4A). SerpinB5 has demonstrated strong antithrombotic and anticoagulant activity in both in vivo and in vitro studies. It may prolong prothrombin time (PT) and activated partial thromboplastin time (APTT) and inhibit thrombosis in the *in vivo* models[26]. If compensatory effects are present, we hypothesize that AaaaNAT1 may interact with these proteins or possess similar anticoagulant properties. Initially, we expressed the AaaaNAT1 protein in vitro (Fig. 4B) for subsequent anticoagulation testing. The blood coagulation experiment showed that, using PTYB empty protein as the control, AaaaNAT1 at an equivalent concentration significantly prolonged blood coagulation time (Fig. 4C), comparable to the effect of sodium citrate. Furthermore, AaaaNAT1 may substantially inhibit platelet aggregation (Fig. 4D). We categorized AaaaNAT1 into various concentration groups and evaluated four coagulation parameters. The results showed that AaaaNAT1 markedly extended APTT (Fig. 4E), PT (Fig. 4F), TT (Fig. 4G), and FIB (Fig. 4H), with the extension increasing at higher aaNAT1 concentrations. Thus, current data suggest that *AaaaNAT1* affects host platelet aggregation and blood coagulation. It is a powerful anticoagulant protein located in the salivary glands of *Aedes aegypti* mosquitoes, essential for their blood-feeding activity.

**Fig. 4.**
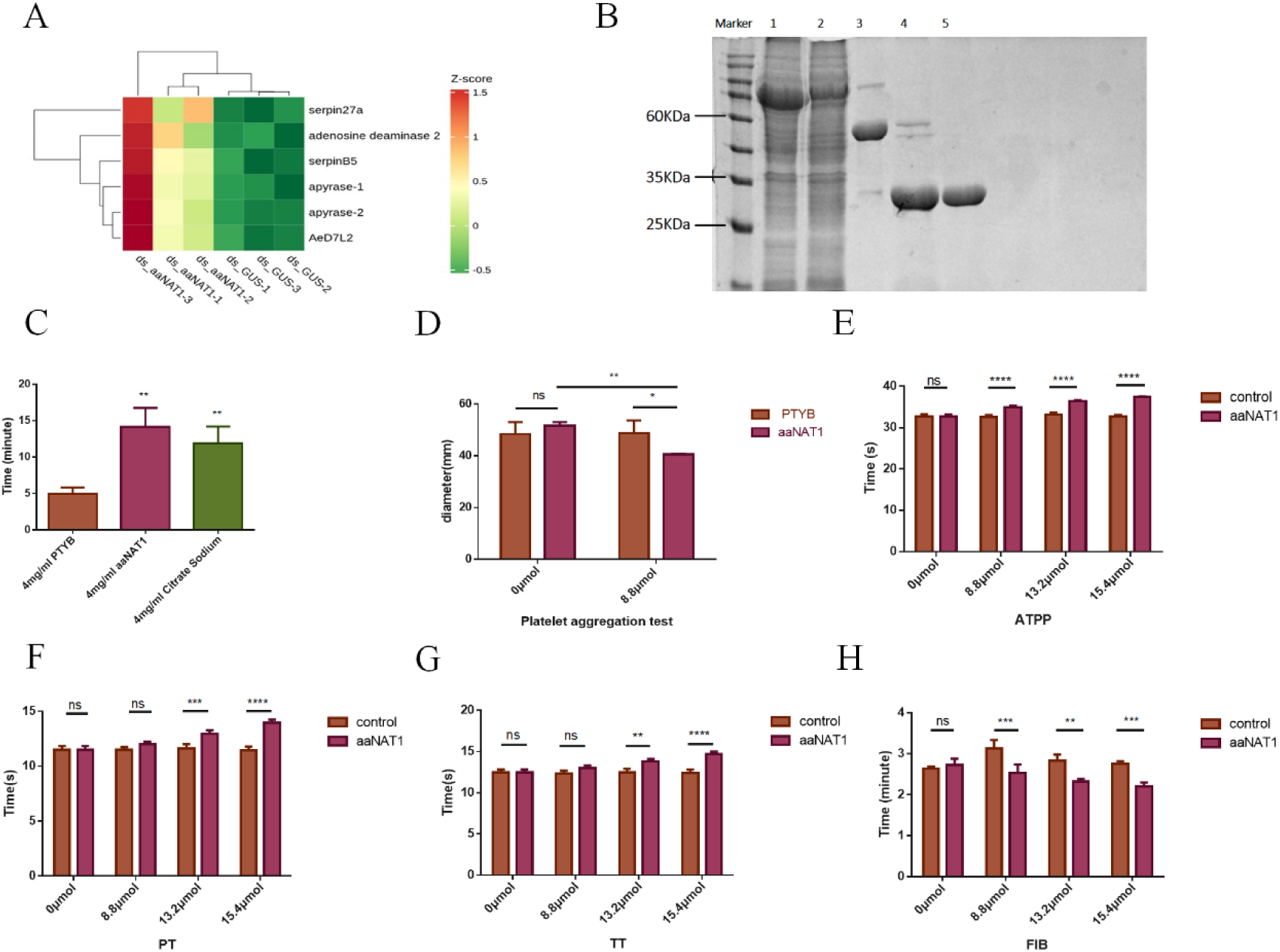
Heatmap of coagulation-related substances and anticoagulation function verification chart. (**A**) Heatmap of proteins with significant changes in proteome function affecting blood coagulation. (**B**) Confirmation of in vitro expression of the aaNAT1 protein (1: Supernatant; 2: FT; 3: R; 4–5: Protein). (**C**) The blood coagulation time in the aaNAT1 protein group and the positive control group (sodium citrate), using an identical concentration of PTYB empty protein as control, was significantly prolonged compared to the control group. (**D**) Using PTYB empty protein at an equivalent concentration as a control, and establishing two concentration groups (0 μM and 8.8 μM), platelet aggregation decreased with increasing aaNAT1 protein concentration. AaaaNAT1 protein was prepared at four concentrations: 0 μmol, 8.8 μmol, 13.2 μmol, and 15.4 μmol, with an equivalent concentration of PTYB empty vector protein as control. (E–H) The results of four coagulation assays showed that prothrombin activation times in (**E**) APTT, (**F**) PT, (**G**) TT, and (**H**) FIB tests were significantly prolonged compared to the control group. \*\*\*\**p* < 0.0001, \*\*\**p* < 0.001, \*\**p* < 0.01, \**p* < 0.05.

### 4. *AaaaNAT1* extends the host’s coagulation time by eliminating norepinephrine from the host

AaaaNAT1 exhibits broad substrate specificity, and its known substrates include serotonin, norepinephrine, and other neurotransmitters[4], which play important roles in physiological processes. Moreover, serotonin and norepinephrine are key regulators of platelet aggregation and blood coagulation, promoting platelet clumping and accelerating coagulation [27]. To identify the specific substances *AaaaNAT1* interacts with to exert its anticoagulant effect, we examined metabolomic results. Metabolomic analysis revealed that among AaaaNAT1’s substrates, only norepinephrine and octopamine levels changed significantly—and interestingly, in opposite directions— while levels of other neurotransmitter substrates remained unchanged (Fig. 5). Currently, there is no direct evidence linking octopamine to blood coagulation, but norepinephrine has been shown to bind adrenergic receptors on platelet surfaces, activating signaling pathways that promote platelet activation and aggregation. Here, metabolomic results showed that norepinephrine levels increased when *AaaaNAT1* expression was downregulated. It is reasonable to hypothesize that AaaaNAT1, as a salivary protein, preferentially acetylates norepinephrine to slow platelet activation and aggregation, thereby facilitating mosquito blood feeding. To confirm this hypothesis, we first used ITC to test the affinity of AaaaNAT1 for different substrates. The results showed that AaaaNAT1 has the strongest binding affinity to norepinephrine compared with dopamine and serotonin (Table 1 and Fig. 6). This suggests that AaaaNAT1 may preferentially respond to norepinephrine in the presence of multiple substrates in the tissue environment. Subsequently, we dissected and collected the salivary glands of *Aedes aegypti* after knocking down *AaaaNAT1* and extracted the total salivary gland protein. After injection into the hands of mice, ELISA results showed that norepinephrine levels in the hands were significantly increased in the *AaaaNAT1* knockdown salivary protein group compared with the wild-type group (Fig. 7A).

**Fig. 5.**
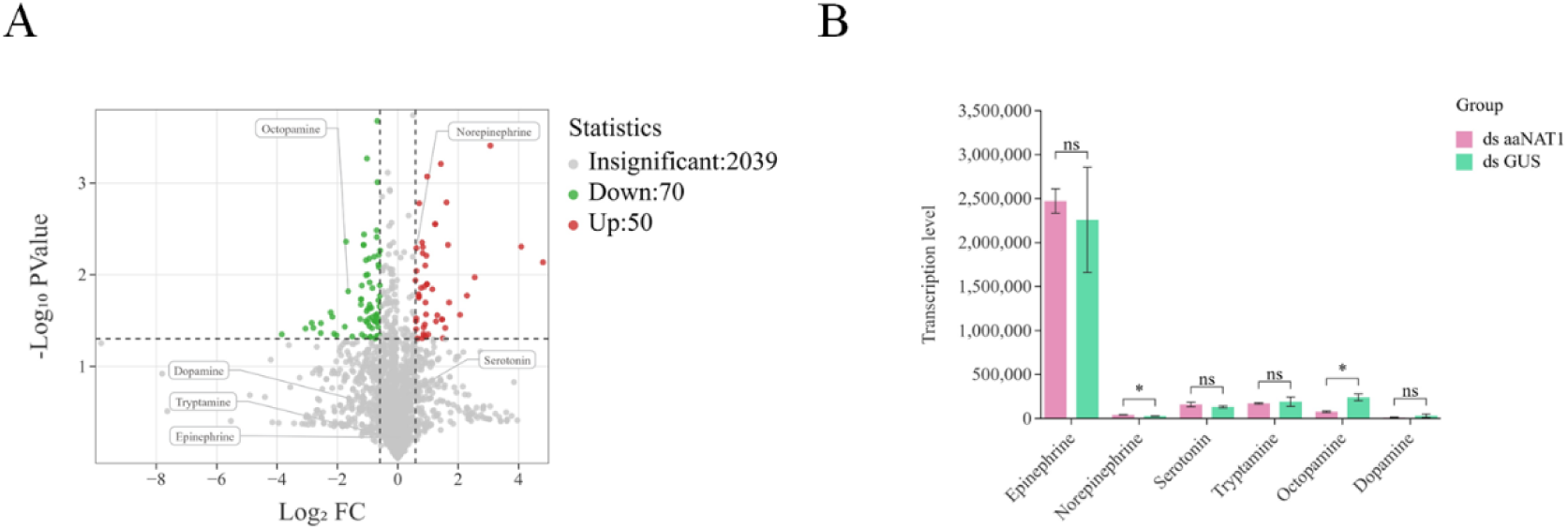
Substrate changes after knocking down *AaaaNATI*. (**A**) Volcano plot of differential metabolites, highlighting substrates of *AaaaNAT1* detected in the metabolome. (**B**) Bar graph of changes in substrate content after knockdown of *AaaaNAT1*. *ds aaNAT1*: the group fed with *aaNAT1* dsRNA; *ds GUS*: the group fed with *GUS* dsRNA. \**p* < 0.05.

**Table 1.** Thermodynamic parameters of aaNAT1 protein by ITC.

| Table 1 Thermodynamic parameters of aaNAT1 protein by ITC. |  |  |  |  |  |
| --- | --- | --- | --- | --- | --- |
| Ligand | Binding | Stoichiometry | $\Delta H$, Kcal/mol±SE | -T$\Delta S$, kcal/mol | KD, M |
| Norepinephrine | Yes | 3.23±0.179 | -46.2±6.45 | 40.0 | 28.5e-6 |
| Dopamine | Yes | 8.64 | -20.7 | 14.6 | 33.2e-6 |
| Serotonin | Yes | 0.153 | -1.00 | -2.62 | 2.22e-3 |

**Fig. 6.**
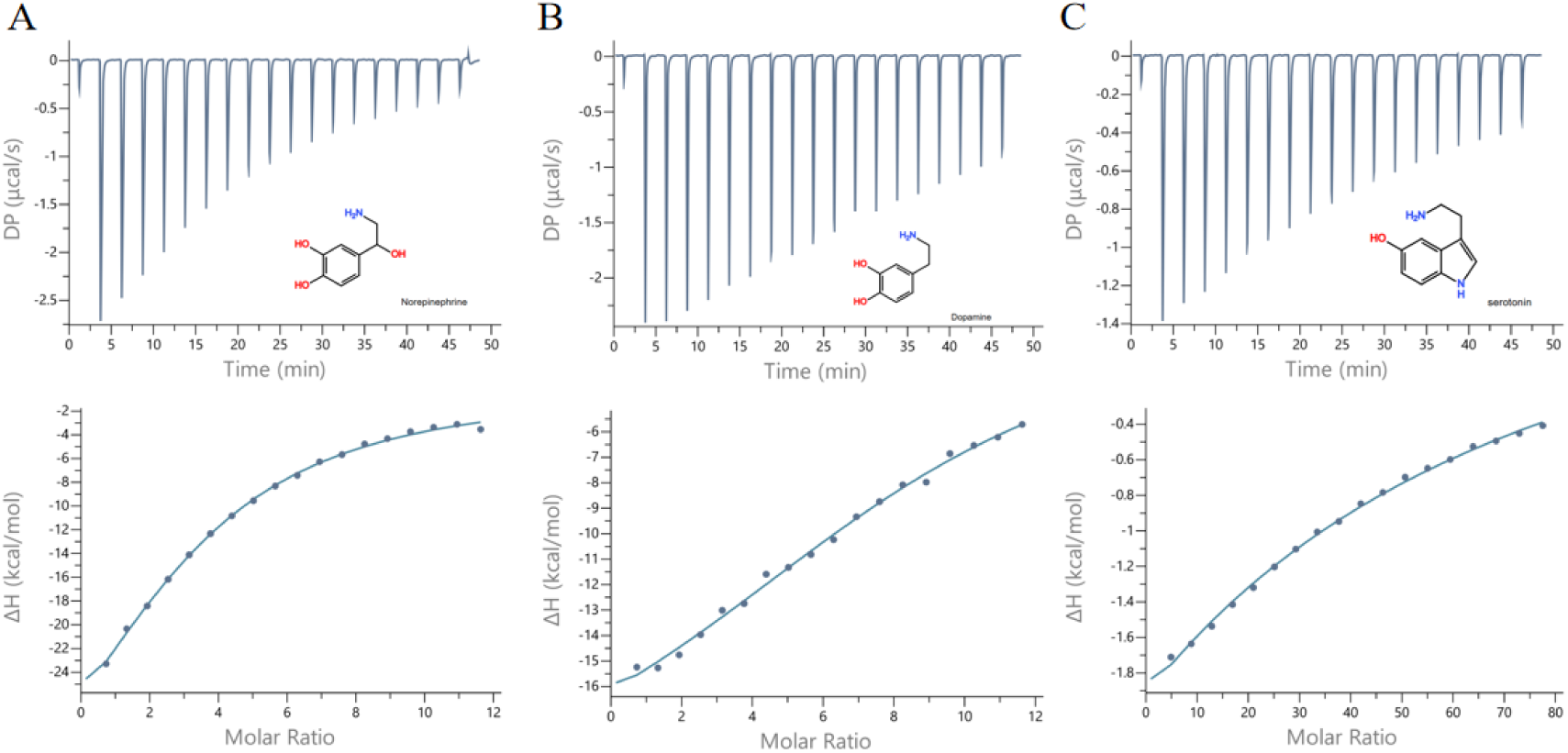
Binding of biogenic amines to aaNAT1 by ITC. In each panel, the upper curve shows the observed heat for each injection. The lower graph shows the enthalpies for each injection and their fit to a single-site binding model used to calculate thermodynamic parameters. Panels: norepinephrine (**A**), dopamine (**B**), serotonin (**C**). Insets show the names and chemical formulas of the compounds.

**Fig. 7.**
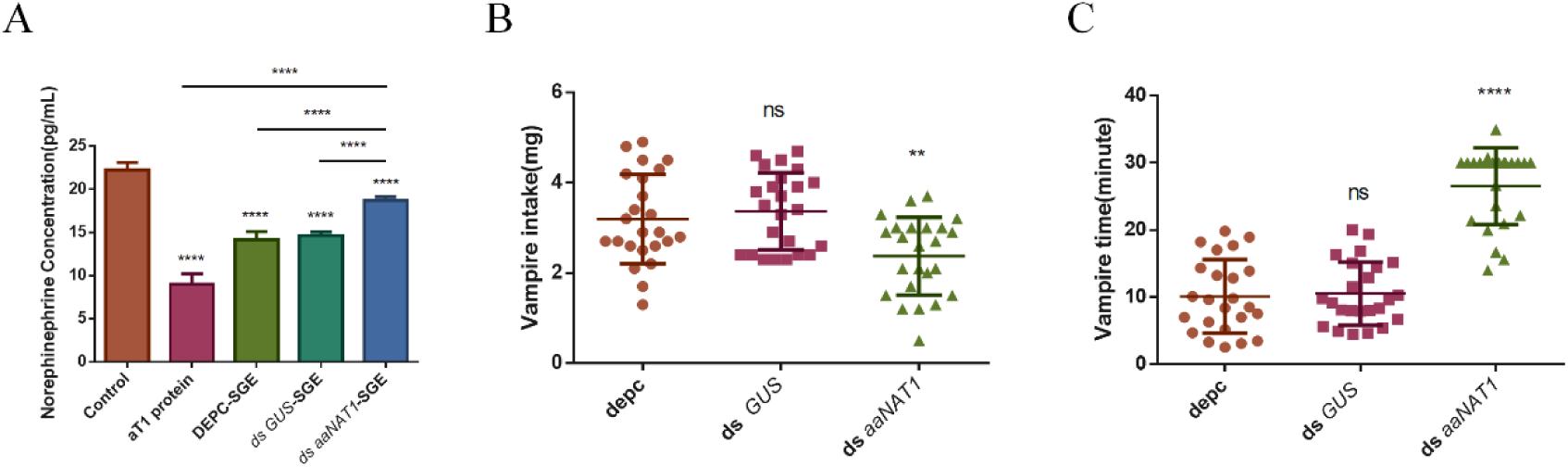
Effects of AaaaNAT1 on norepinephrine content and mosquito blood-sucking behavior. The salivary gland proteins of *Aedes aegypti* from different treatment groups were extracted and injected into the hands of mice. Norepinephrine content in mouse hand tissue was measured by ELISA. The salivary protein group after *AaaaNAT1* knockdown could not effectively remove norepinephrine from mouse tissue (**A**). At the same time, *Aedes aegypti* with *AaaaNAT1* knockdown and the control group were starved for 12 h before blood feeding. The blood intake of the knockdown group was significantly reduced (**B**), and the feeding time was significantly prolonged (**C**). Control: Mouse hand injected with DEPC water; aT1 protein: Mouse hand injected with AaaaNAT1 protein; DEPC-SGE: Mouse hand injected with salivary gland extract from DEPC water-injected mosquitoes; *ds GUS*-SGE: Mouse hand injected with salivary gland extract from GUS dsRNA-injected mosquitoes; *ds aaNAT1*-SGE: Mouse hand injected with salivary gland extract from *AaaaNAT1* dsRNA-injected mosquitoes; *ds GUS*: Mosquitoes injected with GUS dsRNA; *ds aaNAT1*: Mosquitoes injected with *AaaaNAT1* dsRNA; DEPC: Mosquitoes injected with DEPC water. \*\*\*\**p* < 0.0001, \*\**p* < 0.01.

**Fig. 8.**
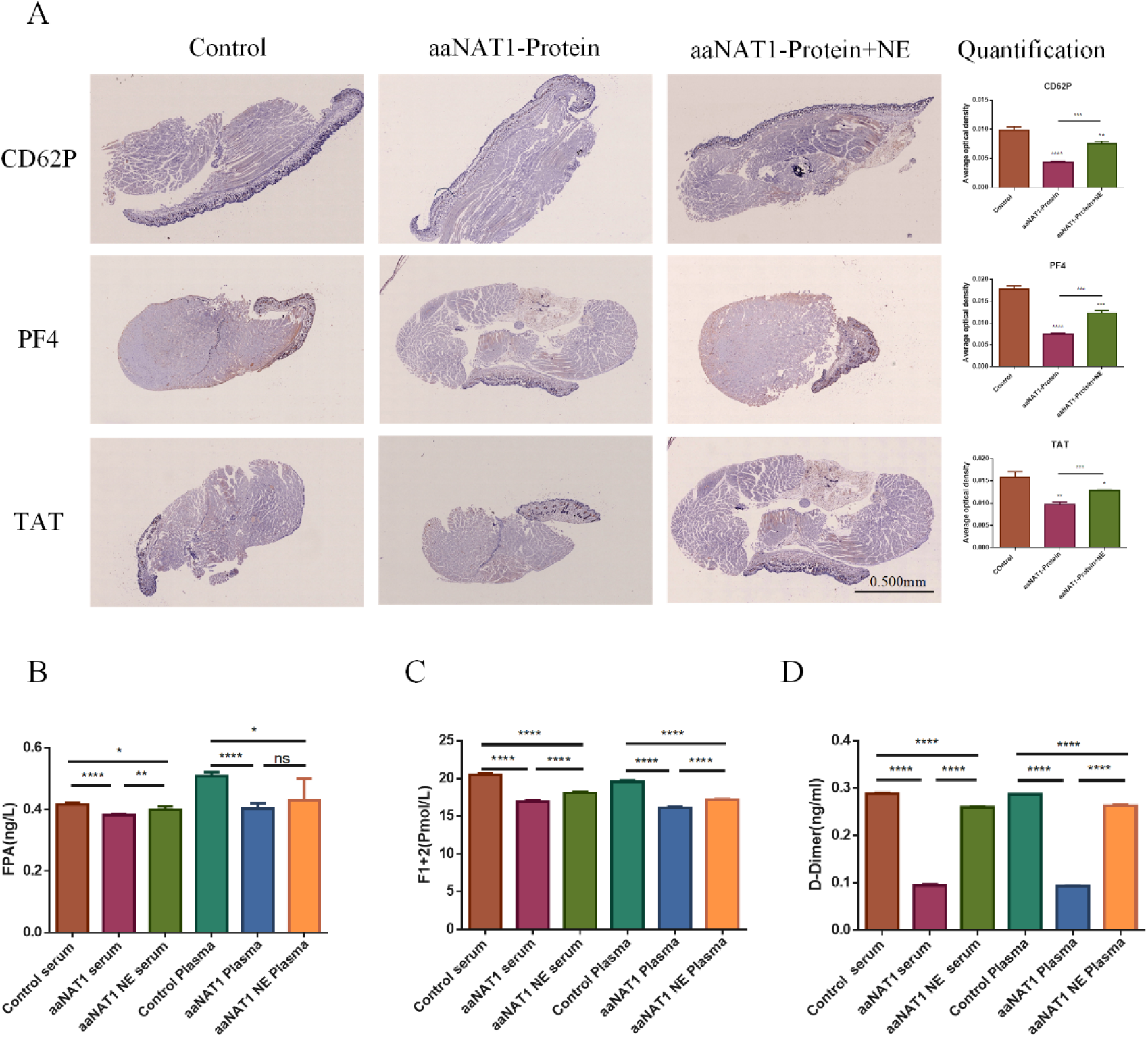
Immunohistochemistry and ELISA. CD62P, PF4, and TAT antibodies were used to stain the legs of mice injected with DEPC water, AaaaNAT1 protein, or a mixture of AaaaNAT1 protein and norepinephrine. Brown particles indicate positive expression. The left side of the figure indicates the antibodies used, the top indicates the different treatment groups, and the right shows the quantitative results of immunohistochemical staining. Control groups: Control: Mouse leg injected with DEPC water; aaNAT1 protein: Mouse leg injected with AaaaNAT1 protein; aaNAT1 protein + NE: Mouse leg injected with a mixture of AaaaNAT1 protein and norepinephrine. Scale: 1:30 (**A**) ELISA was used to detect the levels of (**B**) FPA, (**C**) F1+2, and (D) D-Dimer in serum and plasma from different treatment groups. Control Serum: Serum of mice injected with DEPC water; Control plasma: plasma of mice injected with DEPC water; aaNAT1 Serum: Serum of mice injected with aaNAT1 protein; aaNAT1 plasma: plasma of mice injected with aaNAT1 protein; aaNAT1 NE Serum: Serum of mice injected with the mixture of aaNAT1 protein and norepinephrine; aaNAT1 NE protein; aaNAT1 plasma: plasma of mice injected with the mixture of aaNAT1 protein and norepinephrine. \*\*\*\**p* < 0.0001, \*\*\**p* < 0.001, \*\**p* < 0.01, \**p* < 0.05.

Furthermore, we observed a significant reduction in blood intake by *Aedes aegypti* after *AaaaNAT1* knockdown (Fig. 7B), along with an increase in feeding time (Fig. 7C), indicating that *AaaaNAT1* significantly affects mosquito blood-feeding behavior. Previous results of this study also showed that AaaaNAT1 slowed the activation of the coagulation cascade and fibrinolysis. It is well known that platelet aggregation, the coagulation cascade, and the fibrinolytic system are interrelated. For example, when platelet aggregation is blocked, substances such as t-PA released by platelets are reduced, resulting in decreased plasminogen activation and impaired fibrinolysis [28]. Similarly, when platelet aggregation is blocked, activation of coagulation factors on the platelet surface and thrombin complex formation are reduced, leading to lower thrombin production and suppression of the coagulation cascade [29]. Therefore, we also aimed to determine whether AaaaNAT1’s effects on the coagulation and fibrinolytic responses and the coagulation cascade shown by the four results of *AaaaNAT1* coagulation are mediated through its interaction with norepinephrine or through other mechanisms. To further explore this, we injected AaaaNAT1 protein into mice as the experimental group, and AaaaNAT1 protein with norepinephrine as the complementary group. Immunohistochemistry was used to detect P-selectin (CD62P), platelet factor 4 (PF4), and thrombin–antithrombin III complex (TAT). ELISA was used to measure prothrombin fragment (F1+2) in serum and plasma. Fibrin peptide A (FPA) and D-dimer levels were measured to assess activation of platelets, the coagulation cascade, and the fibrinolytic system. The results showed that *AaaaNAT1* significantly inhibited the activation of all three pathways, but this inhibition was markedly relieved with norepinephrine supplementation (Figure). Therefore, we suggest that AaaaNAT1 removes norepinephrine at the wound site during mosquito feeding, thereby reducing host platelet aggregation and inhibiting activation of the coagulation cascade and fibrinolytic system, ultimately slowing host coagulation and enhancing mosquito blood-feeding behavior.

### 5. AaaaNAT1 influences mosquitoes’ host-seeking behavior by modulating octopamine levels in the organism

Octopamine is recognized for its significant influence on mosquito behavior, with octopamine receptors co-expressed in olfactory neurons functioning as chemoreceptors[30]. Olfactory capabilities in insects are modulated by the levels of octopamine and its receptors. Host location in insects relies on multiple senses, among which olfaction is one of the most critical[31,32]. Notably, octopamine was found to be considerably down-regulated in the metabolomic findings. In most studies, the knockdown or deletion of *aaNAT1* elevates its substrate, but octopamine exhibits an atypical response. Furthermore, prior findings indicated that *AaaaNAT1* knockdown in female *Aedes aegypti* mosquitoes resulted in ∼50% of the knockdown strains successfully aspirating blood even after 12 h of starvation, though host-seeking duration was prolonged (Fig. 7C). Given the role of octopamine and the experimental findings, it was questioned whether this behavioral change was due to its down-regulation. To determine if the altered behavior was linked to reduced octopamine levels or impaired olfaction/audition, subsequent experiments were conducted. First, octopamine receptor levels were assessed after *AaaaNAT1* knockdown and subsequent restoration, while blood-feeding behavior reversals were monitored. Through homology comparison with Drosophila melanogaster, four octopamine receptors were identified in Aedes aegypti, with expression levels varying in relation to blood feeding, linking them to feeding behavior[33]. These receptors are octopamine receptor α-1(OAα-1R), octopamine receptor α-2(OAα-2R), octopamine receptor β-2(OAβ-2R) and octopamine receptor β-3(OAβ-3R). Their expression and octopamine levels were measured at different time points post-*AaaaNAT1* knockdown. All four receptors were upregulated at 2 h post-*AaaaNAT1* knockdown but significantly downregulated by 12 hours (Fig. S3). AaaaNAT1 knockdown caused a sustained reduction in octopamine levels. Co-injection of *AaaaNAT1* dsRNA with 10 mM octopamine restored receptor mRNA levels and octopamine concentrations to 70–80% of control levels (Fig. 9A-E). Comparisons were made among the blank control, negative control, experimental (*AaaaNAT1* dsRNA), and supplementation (*AaaaNAT1* dsRNA + octopamine) groups. Host-seeking duration was markedly prolonged in the experimental group compared to the two control groups, while the supplemented group showed intermediate results (Fig. 9F). These findings suggest that AaaaNAT1 may influence *Aedes aegypti* host-seeking behavior by regulating octopamine levels and affecting the octopamine-octopamine receptor signaling pathway levels.

**Fig. 9.**
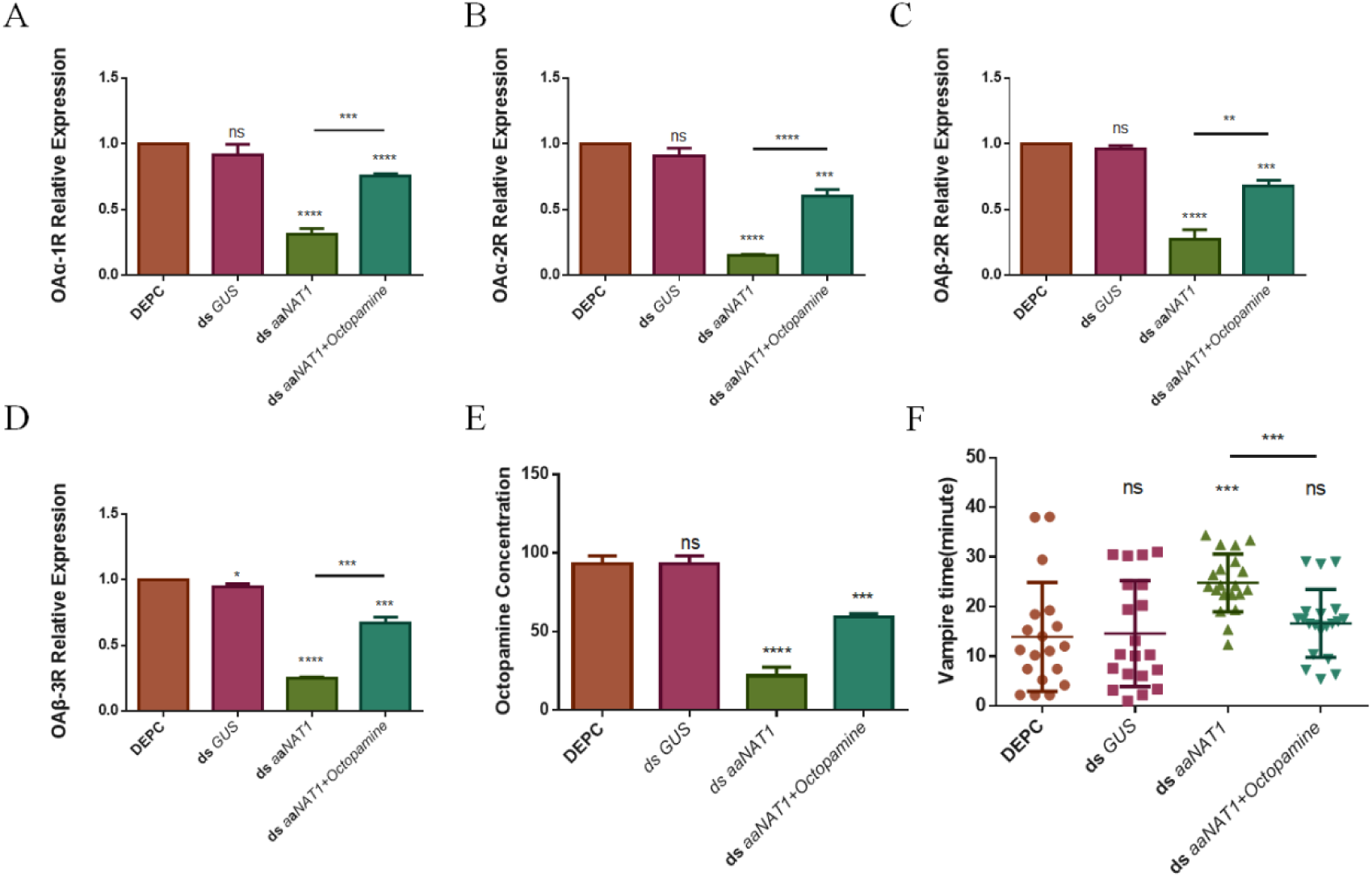
Octopamine receptor expression and octopamine content after different treatments. Detected the OAα-1R(**A**), OAα-2R(**B**), OAβ-2R(**C**) and OAβ-3R(**D**) levels in *Aedes aegypti* across several treatment groups were quantified using qPCR, while the receptor content in the body was simultaneously assessed via ELISA(**E**), alongside observations of alterations in the mosquitoes’ blood-feeding behavior(**F**). DEPC: *Aedes aegypti* injected with DEPC water; *ds GUS*: *Aedes aegypti* injected with the GUS dsRNA; *ds aaNAT1*: *Aedes aegypti* injected with the *AaaaNAT1* ds RNA; *ds aaNAT1*: Octopamine: *Aedes aegypti* injected with the mixture of *AaaaNAT1* ds RNA and octopamine. \*\*\*\**p* <0.0001, \*\*\**p*<0.001 \*\**p*<0.01, \**p*<0.05.

## Discussion

The enzyme aaNAT was first identified and extensively studied in vertebrates, where it acetylates serotonin to produce melatonin [34–36]. This process is critical—and possibly essential—for melatonin biosynthesis. As a result, aaNAT, often termed a “chronoenzyme,” links melatonin production to the regulation of circadian rhythms. Studies in fish species such as *Esox lucius* [37], *Oncorhynchus mykiss* [38,39], *Danio rerio* [40], and *Sparus aurata* [41] typically reveal two types of aaNATs: aaNAT1 and aaNAT2. In contrast, only a single *aaNAT* gene has been identified in mammals and birds [35], where its role in circadian regulation is more emphasized. Notably, insects possess a higher number of *aaNAT* or *aaNAT-like* genes than vertebrates and mammals[4,8,13,42]. These genes display distinct expression patterns across tissues and developmental stages, suggesting diverse roles in insect biology[43–47]. In *Aedes aegypti*, 13 aaNAT proteins were identified[4,9]. aaNAT1, in particular, has been implicated in melatonin synthesis, sleep regulation, and skin pigmentation across various organisms [9]. In *Bactrocera dorsalis*, *aaNAT1* knockout led to the accumulation of several monoamines in the midgut, altered ROS levels, and reduced expression of *CncC* and *Maf*, which in turn affected GST and P450 detoxifying enzyme activity [18]. These findings indicate that beyond its traditional role in melanization, aaNAT1 also contributes to detoxification processes in insects.

This study revealed a novel anticoagulant function of aaNAT1, distinct from previously known roles. The mechanism by which aaNAT1 acts as an anticoagulant has been largely elucidated. Blood-feeding directly affects mosquito ovarian maturation and oviposition[3,30,48], and is therefore critical to the transmission of yellow fever viruses[1,49]. Anticoagulant proteins in the salivary glands play a key role in this process[3]. If potent anticoagulant proteins can influence mosquito blood-feeding behavior, they could not only curb virus transmission but also help control mosquito populations. Host coagulation occurs in three phases: vasoconstriction[2], platelet aggregation[20,50], and blood coagulation. Vasoconstriction reduces blood flow to the wound site, platelets form an initial plug, and fibrin reinforces this plug to achieve hemostasis and support wound healing[50]. This study not only confirmed the anticoagulant activity of aaNAT1 but also showed that it modulates host platelet aggregation and mosquito host-seeking behavior by regulating norepinephrine and octopamine levels. These findings deepen our understanding of aaNAT1 in *Aedes aegypti* and identify it as a potential target for mosquito control. They also suggest a new direction for developing antithrombotic drugs, as anticoagulant proteins have shown promise in antithrombotic therapy[51].

This study employed multiple experimental methodologies to clarify the function of *Aedes aegypti*’s aaNAT1 in anticoagulation. Initially, an *AaaaNAT1* knockdown strain was generated using RNA interference, and several pairs of salivary glands were harvested for transcriptomic, metabolomic, and proteomic analyses. The integration of in vivo and in vitro investigations indicates that AaaaNAT1 downregulation directly leads to norepinephrine accumulation; thus, AaaaNAT1 likely facilitates *Aedes aegypti* blood feeding by lowering norepinephrine levels near the host wound and prolonging platelet aggregation time. Integrating the proteomics data with coagulation assays revealed that AaaaNAT1 not only influenced platelet aggregation but also significantly delayed coagulation factor activation. Inhibiting the activation of coagulation factors— the most rapid and effective mechanism of blood clotting—creates highly favorable conditions for mosquito blood feeding. In addition to its primary role in blood coagulation, *AaaaNAT1* knockdown reduced blood intake and extended the host-seeking duration of *Aedes aegypti*, potentially linked to octopamine content. The interaction between octopamine and its receptors may affect the mosquito’s auditory and olfactory abilities, which are critical for host detection. Downregulation of octopamine renders mosquitoes less responsive to host cues, impairing their detection ability and resulting in prolonged host-seeking behavior following *AaaaNAT1* interference. Nevertheless, several questions require further investigation. We confirmed that *AaaaNAT1* ablation affected the mRNA levels of four octopamine receptors and altered octopamine concentration in *Aedes aegypti*. Behavioral experiments verified the correlation between the time taken by mosquitoes to find a host and the expression levels of octopamine receptors and octopamine concentration. How does AaaaNAT1 modulate octopamine concentrations in *Aedes aegypti*? Preliminary omics analysis suggests that changes in the regulation of multiple genes and metabolites in the tryptophan and tyrosine metabolic pathways may influence octopamine synthesis; however, the mechanisms of regulation remain unclear and warrant further study.

The current findings enhance our understanding of the molecular mechanisms underlying mosquito blood feeding and highlight the potential of AaaaNAT1 as a target for disrupting mosquito feeding and transmission of mosquito-borne diseases. Altering AaaaNAT1 levels can impair mosquitoes’ ability to detect hosts and acquire blood, offering a novel strategy for vector control. Moreover, these findings underscore the complexity of mosquito physiological adaptation to blood feeding and the intricate interplay among its neurological, endocrine, and biochemical systems. The regulation of the biogenic amine octopamine by AaaaNAT1 suggests a link between neurotransmission and mosquitoes’ capacity to locate and feed on hosts, emphasizing the dynamic nature of mosquito-host interactions. This understanding may support the development of innovative strategies to reduce the impact of mosquito-borne diseases. Given the strong anticoagulant activity of aaNAT1, it may also offer a new avenue for antithrombotic drug development and advance progress in human medicine.

## Conclusion

This study elucidates a novel and critical role of Arylalkylamine N-acetyltransferase-1 (aaNAT1) in Aedes aegypti, demonstrating its dual function in modulating host coagulation pathways and mosquito blood-feeding behavior. Through multi-omics analyses combined with biochemical and behavioral assays, we revealed that aaNAT1 exerts its anticoagulant effects by preferentially acetylating norepinephrine at the host wound site, thereby inhibiting platelet aggregation and delaying the coagulation cascade and fibrinolysis. Furthermore, aaNAT1 knockdown disrupted octopamine signaling, impairing host-seeking efficiency and reducing blood-feeding success, highlighting its broader physiological impact beyond traditional roles in pigmentation and immunity.

## Materials and Methods

### 1. Animal

*Aedes aegypti* larvae were raised on an artificial diet under controlled conditions (16h light: 8h dark; temperature 26 ± 1 °C; humidity 60–70%). Larvae were fed a starch-based artificial diet, followed by 8% sugar water after adult emergence. Female mosquitoes three days post-eclosion were selected for subsequent experiments.

The Kunming mice utilized in this experiment were supplied by Jiangsu Huachuang sino Pharma Tech Co.,Ltd. and were maintained in the animal facility of Hainan University, which maintained a temperature of 22 ± 2℃ and a 12-hour light-dark cycle. All animal experimentation methods received approval from the Animal Ethics Committee of Hainan University (Ethical Number: HNUAUCC-2025-00482).

### 2. *aaNAT1* dsRNA expression

Female mosquito cDNA was used as the template, with aaNAT1-F (GCGGCCGCGGTTGTGAGCATCCCAAGTT) and aaNAT1-R (AACCGCAGT GTGTGGTTTCTCTCGAG) primers used to amplify the PCR product. The fragment was ligated into the expression vector PL4440 and transformed into HT115 competent cells for dsRNA expression. The negative control, *β*-*glucuronidase* (GUS) gene (KY848224), was obtained from original laboratory strains [52].

### 3. RNAi and salivary gland collection

PL4440-aaNAT1 and PL4440-GUS bacterial solutions were resuscitated, added to 1 L LB liquid medium, and incubated for 4 h. IPTG was added, and expression was induced for 5h at 37°C. The adult mosquitoes, three days post-eclosion, were interfered with using a previously reported method [53]. Female mosquitoes, eight days after interference, were anesthetized at −20 °C, their salivary glands were dissected in normal saline, rapidly transferred to dry ice, and stored at -80 °C. A total of 5,000 pairs of female mosquito salivary glands were collected from each group.

### 4. Proteome

The salivary glands of female mosquitoes were collected after 8 days of continuous interference, with 3,000 pairs collected per group (*ds*-*aaNAT1* group, n = 3; *ds*-*GUS* group, n = 3). Protein extraction, detection, and quantitative analysis were performed by Wuhan Metware Biotechnology Co., Ltd. (www.metware.cn). A threshold of *p* ≤ 0.05 was used to screen differentially expressed proteins. Kyoto Encyclopedia of Genes and Genomes (KEGG) (https://www.genome.jp/kegg/), Gene Ontology (GO) (http://geneontology.org/), Cluster of Orthologous Groups of proteins (COG) (http://www.ncbi.nlm.nih.gov/COG) were used for protein function annotation and pathway enrichment analysis. InterPro (https://www.ebi.ac.uk/interpro/) was used for protein domain structure analysis. GO and KEGG enrichment analyses of differentially expressed proteins were performed using ClusterProfiler with a threshold of *p* ≤ 0.05. The log2 fold change and the absolute log10 *p*-value were used to generate a volcano plot. Unsupervised PCA (principal component analysis) was performed using the prcomp function in R (www.r-project.org).

### 5. Metabolome

The salivary glands of female mosquitoes were collected after 8 days of continuous interference, with 2000 pairs collected per group (*ds aaNAT1* group, n = 3; *ds GUS* group, n = 3). Metabolite extraction, detection, and quantitative analysis were performed by Wuhan Metware Biotechnology Co., Ltd. (www.metware.cn). Unsupervised PCA was conducted using the prcomp function in R (www.r-project.org) after unit variance scaling of the data. The log2 of the fold change and the absolute log10of the p-value for each differential metabolite were calculated to generate a volcano plot. HCA results for samples and metabolites were presented as heatmaps with dendrograms, while Pearson correlation coefficients (PCC) between samples were calculated using the cor function in R and displayed as heatmaps. Both HCA and PCC analyses were performed using the R package Complex Heatmap. For HCA, normalized metabolite signal intensities (unit variance scaled) were visualized as a color spectrum. For two-group comparisons, differential metabolites were identified based on VIP > 1 and *p* < 0.05 (Student’s t-test). Finally, identified metabolites were annotated using the KEGG Compound database (http://www.kegg.jp/kegg/compound/), and mapped to the KEGG Pathway database (http://www.kegg.jp/kegg/pathway.html). Pathways with significantly regulated metabolites were subjected to MSEA (metabolite set enrichment analysis), with significance determined by hypergeometric test p-values.

### 6. Transcriptome sequencing

Salivary glands of female mosquitoes after 8 days of continuous interference were collected (2000 pairs per group; *ds aaNAT1* group, n = 3; *ds GUS* group, n = 3), snap-frozen on dry ice, and stored at -80 °C. RNA library preparation and RNA-seq were performed by Wuhan Metware Biotechnology Co., Ltd. RNA sequencing was conducted on the Illumina platform. Differential expression analysis was performed using DESeq2, with thresholds of |log2Fold Change| ≥1 and p < 0.05. Differentially expressed genes were standardized using the scale function in R, followed by K-means clustering. GO and KEGG pathway enrichment analyses were conducted using clusterProfiler (v4.6.0). A volcano plot was generated using the log2fold change and the absolute log10 p-value for each differential gene. Unsupervised PCA was performed using the prcomp function in R (www.r-project.org).

### 7. Quantitative real-time PCR analysis

A total of nine genes showing significant changes in transcriptomic analysis were selected for data validation. The primer sequences for aaNAT1, RPS17 and the nine selected genes are listed in Table S1. Quantitative primers for the four octopamine receptors were obtained from the literature[33]. Total RNA was extracted from mosquitoes using the Trizol method, and cDNA was synthesized using the SPARKscript II RT Plus Kit (with gDNA Eraser, Sparkjade). The reaction mixture was prepared according to the instructions of the 2×SYBR Green qPCR Mix (with Rox, Sparkjade) and added to a 96-well plate. qPCR was performed using a LightCycler96 instrument (Roche). RPS17 was used as an internal control to normalize cDNA concentration differences among samples. Gene expression levels were calculated using the 2^-ΔΔCt^ method, and statistical analysis was conducted using a t-test in GraphPad Prism 6.02.

### 8. Cloning, expression, and purification of recombinant proteins

PCR products were amplified using primers and mosquito cDNA as templates. The recombinant plasmid was cloned into the PTYB vector and transformed into *Eschericihia coli* DH5α. After sequencing, the plasmid was transformed into *E. coli* BL21 (DE3), and the strain was preserved. A 3L culture was incubated at 37°C for 3-4 h, followed by induction with IPTG at 16°C for 24h. After centrifugation, the bacterial pellet was lysed using an ultrasonic processor at 70% efficiency (3 s ON, 3 s OFF) for 15 min until the lysate became clear. The supernatant containing the expressed proteins was collected after centrifugation at 8000×g for 30 min at 4°C. Proteins were purified using chitin beads and cleaved in β-mercaptoethanol at 23°C for 40 h. The eluted solution contained the purified protein, which was concentrated using an Amicon Ultra-0.5 column, Protein concentration was measured using a BCA assay kit (Beyotime) and a microplate reader. The final protein product was stored at -80°C.

### 9. Isothermal titration calorimetry (ITC)

Thermodynamic binding parameters of aaNAT1 with dopamine, serotonin, and norepinephrine were determined using a MicroCal PEAQ-ITC microcalorimeter. Ligand and acetyl-CoA solutions (600 µM) and protein solution (10 µM) were prepared in pure water. A 5 µL mixture of ligand and acetyl-CoA was titrated into the protein sample in the calorimeter cell at 300 s intervals. The experiments were conducted at 25°C, and thermodynamic parameters were analyzed using MicroCal PEAQ-ITC Analysis Software (V1.41).

### 10. Platelet aggregation assay

The gathered blood was first mixed with sodium citrate anticoagulant at a 9:1 ratio. Next, 380μL of this mixture was combined with 50 μL of recombinant protein. After thorough mixing, the solution was transferred to a sample plate containing 20 μL of CaCl_2_and immediately analyzed using thromboelastography (CFMS LEPU-8800, Beijing Lepu Medical Technology Co., Ltd.). The same concentration of PTYB empty protein served as the control. The experiment was repeated three times, and statistical analysis was performed using Two-Way ANOVA and t-test in GraphPad Prism 6.02.

### 11. Blood clotting tetrachoric

The venous blood of laboratory-raised New Zealand white rabbits was mixed with sodium citrate anticoagulant at a 1:9 ratio. The samples were immediately centrifuged at 3000 rpm for 15 min. After centrifugation, the upper plasma layer was carefully aspirated using a pipetting gun. Four coagulation tests were performed using an automatic biochemical analyzer and corresponding coagulation test discs (Tianjin Micro and Nano Core Technology Co., LTD.). The experiment was divided into four groups, each containing 100 µL of aaNAT1 protein at concentrations of 0 µmol, 8.8 µmol, 13.2 µmol, and 15.4 µmol, respectively, with PTYB no-load protein as the control. The experiment was repeated three times, and statistical analysis was conducted using Two-Way ANOVA and t-test in GraphPad Prism 6.02.

### 12. In vitro transcription of dsRNA and injection

For *aaNAT1* dsRNA synthesis, mosquito cDNA was used as a template, and PCR was performed using primers (F:GGATCCTAATACGACTCACTATAGGGCCGAGGATGTCCTGAAATTG/R:G GATCCTAATACGACTCACTATAGGGCCGAGGATGTCCTGAAATTG), which included the T7 promoter sequence. The *GUS* gene was used as a control. Its PCR product was amplified from the lab-stored GUS-PL440 [52] strain using primers (FGGATCCTAATACGACTCACTATAGGGATGAACATGGCATCGT GGT/R: GGATCCTAATACGACTCACTATAGGGGGCACAGCACATCAAAGAGA) for GUS dsRNA synthesis. Both PCR products were purified, and the corresponding dsRNAs were synthesized using the T7 RiboMAX^TM^ Express RNAi System (Promega). For co-injection experiments, equal volumes of *AaaaNAT1* ds RNA and 10mM octopamine (prepared from octopamine hydrochloride, MedChemExpress (MCE), using deionized water) were mixed. Three-day-old female mosquitoes were injected with 1μL of the mixture. Each treatment group consisted of 100 mosquitoes, and the experiment was repeated three times. qPCR was used to evaluate RNA interference efficiency.

### 13. Measuring the survival rate, time and volume of blood aspiration

Post-injection, mosquito awakening time was recorded, and no food was provided. Mortality was recorded after 12 hours. Blood-feeding behavior was assessed by placing mosquitoes on anesthetized mice fixed at the bottom of the cage. Mosquitoes were weighed before and after feeding to evaluate blood intake. Blood-feeding was initiated when mosquitoes contacted the mouse and ended when feeding stopped. The data collected from weight changes and feeding duration are then used to statistically analyze blood consumption across the mosquito population. Mosquitoes were allowed to feed again in three blood-feeding cages, corresponding to *ds aaNAT1*, *ds GUS*, and DEPC groups. The experiment was repeated three times, and statistical analysis was conducted using a t-test in GraphPad Prism 6.02.

### 14. Determination of octopamine content

*Aedes aegypti* samples were collected at specified time points post-injection and stored on ice. Three sterilized, room-temperature steel balls (3 mm diameter) were added to each tube, and samples were crushed using a freezing crusher for 10 min, followed by centrifugation at 5000 g for 10 min. The supernatant was collected and used for octopamine quantification with an insect octopamine detection kit (Shanghai Enzyme Linked Biotechnology Co., LTD.).

### 15. Determination of norepinephrine content

As per experimental requirements, DEPC water, AaaaNAT1 protein, DEPC-SGE, ds GUS-SGE, and ds aaNAT1-SGE were injected into mouse legs. Approximately 1 g of injected leg tissue was excised and homogenized in 9 mL PBS (pH 7.4). The homogenate was centrifuged at 3000 rpm for 20 min, and the supernatant was collected after centrifugation. The norepinephrine content in the mouse legs was measured in different treatment groups using a mouse norepinephrine detection kit (Shanghai Enzyme Linked Biotechnology Co., LTD.).

### 16. Preparation of salivary gland extracts

Mosquitoes were frozen at -20℃ for 3 min, wings and legs were removed, and the bodies were immersed in 75% ethanol to remove surface grease. They were then placed in saline to wash off residual ethanol. Washed mosquitoes were transferred to saline for salivary gland dissection, and the dissected glands were stored in sterile tubes containing PBS (100 glands per tube) and frozen at -80℃. The frozen salivary glands were placed in an ice box and thoroughly homogenized within 5 min. After vortexing the homogenate, it was centrifuged at 12,000 rpm at 4℃ for 20 min. The supernatant was collected as the salivary gland extract (SGE). The crude protein concentration of the SGE was determined by the BCA method and the extract was stored at -80℃.

### 17. Immunohistochemistry

1. Paraffin embedded sections: Mice administered DEPC water, AaaaNAT1 protein, or a combination of AaaaNAT1 and norepinephrine had their legs amputated and immediately immersed in formaldehyde fixative (10× tissue volume) for 48 hours. The fixed tissues were sequentially immersed in graded ethanol (70%, 80%, 90%, 95%, 100%) for 15 min each. Clearing was performed using xylene to remove residual ethanol. The cleaned tissue was then immersed in pre-melted paraffin at 60 °C, with multiple paraffin changes until xylene was fully removed. Paraffin-embedded mouse tissues were cast into molds and incubated at 60°C for 12 h to eliminate air bubbles. Tissues were positioned in molds and maintained at 60°C for 4 h, with gentle agitation every 2 h. After curing the paraffin surface, molds were removed and immersed in 10 °C water. The tissue blocks were sectioned into 4–5μM slices using a microtome.
2. Deparaffinization and Rehydration of Sections: Paraffin sections were sequentially immersed in xylene I (20 min), xylene II (20 min), xylene III (15 min), anhydrous ethanol I (5 min), anhydrous ethanol II (5 min), 85% ethanol (5 min), and 75% ethanol (5 min), followed by a rinse with distilled water.
3. Antigen Retrieval: Sections were placed in citrate antigen retrieval buffer (pH 6.0) or EDTA antigen retrieval buffer (pH 9.0) and microwaved at medium, medium-high, and high power for 5 min each. Excessive buffer evaporation and tissue drying were avoided. After natural cooling, slides were washed in PBS (pH 7.4) by gentle shaking, three times for 5 min each.
4. Blocking Endogenous Peroxidase: Sections were incubated in 3% hydrogen peroxide at room temperature in the dark for 25 min, followed by three PBS (pH 7.4) washes (5 min each).
5. Serum Blocking: After light drying, tissue boundaries were circled with a histological pen (to prevent antibody runoff). Goat serum was applied to cover the tissue and incubated at room temperature for 15 min.
6. Primary Antibody Application: The blocking solution was gently removed, and primary antibody diluted in PBS was applied. Sections were incubated overnight at 4 °C in a humidified chamber with a small amount of water to prevent evaporation.
7. Secondary Antibody Application: Sections were washed three times in PBS (pH 7.4) for 5 min each. After slight drying, HRP-conjugated secondary antibody (HRP-conjugated), matching the host species of the primary antibody, was applied to cover the tissue and incubated at room temperature for 50 min.
8. DAB Staining: Sections were washed three times in PBS (pH 7.4) (5 min each), then incubated with freshly prepared DAB solution. Staining time was monitored under a microscope. Positive staining appeared brownish-yellow. Staining was terminated with a tap water rinse.
9. Counterstaining of Nuclei: Sections were counterstained with hematoxylin for 5–10 min, rinsed with tap water, differentiated with 1% HCl in alcohol for a few seconds, rinsed again, and blued with 1% ammonia solution for 1 min. A final rinse with tap water was performed.
10. Dehydration and Mounting: Sections were dehydrated in 75% ethanol (5 min), 85% ethanol (5 min), anhydrous ethanol I (5 min), anhydrous ethanol II (5 min), and cleared in xylene I (5 min). After slight drying, sections were mounted with neutral gum.
11. Result Interpretation: Nuclei stained with hematoxylin appeared blue, while DAB-positive expression appeared brownish-yellow.

### 18. Detection of FPA, F1+2and D-Dimer content

According to experimental requirements, mice will be injected with DEPC water, AaaaNAT1 protein, or a mixture of AaaaNAT1 protein and norepinephrine. After injection, blood will be collected from the eye sockets into sterile tubes. Following natural coagulation at room temperature for 20 min, the blood will be centrifuged at 3000 rpm and 4°C for 20 min, and the supernatant will be collected as the serum sample for testing. Additionally, blood from the injected mice will be collected into sterile tubes containing sodium citrate anticoagulant. After thorough mixing for 20 min, the blood will be centrifuged at 3000 rpm and 4°C for 20 min, and the supernatant will be collected as the plasma sample for testing. The above serum and plasma samples will be used to detect the corresponding substances using the mouse FPA detection kit, mouse F1+2 detection kit, and mouse D-Dimer detection kit.

## Acknowledgments

This work was supported by the Hainan Province Science and Technology Special Fund (ZDYF2023XDNY061), the National Natural Science Foundation of China (U22A20363) and the Major Science and Technology Plan of Hainan Province (ZDKJ2021035). Express gratitude to Professor Chen Wenxue for his amendments to this article.

## Supporting information

Additional supporting information may be found online in the Supporting Information section at the end of the article.

**S1 Fig. Quantitative validation of transcriptome data.** Randomly select 9 genes with significant changes for qPCR validation, and the trends in quantitative data are consistent with those in transcriptome data. *ds GUS*: Feeding the *GUS* ds RNA group. *ds aaNAT1*: Feeding the *AaaaNAT1* ds RNA group. (A) qPCR results, \*\*\*\**p* <0.0001, *** *p* <0.001, ** *p* <0.01, * *p* <0.05.(B) Transcriptome data, red indicates upregulation, whereas green signifies downregulation. A richer hue indicates a greater fold shift in regulation.

**S2 Fig. KEGG Enrichment chord plot.** Substantial alterations in genes and their associated pathways, concentrating exclusively on the oxidative phosphorylation pathway and metabolic pathways.

**S3 Fig. Changes in octopamine and its receptor content.** After knocking down *AaaaNAT1* in *Aedes aegypti* using RNAi, qPCR detected that OAα-1R(**A**), OAα-2R(**B**), OAβ-2R(**C**) and OAβ-3R(**D**) were significantly increased two hours after knockdown, but significantly downregulated after four hours, but the content of octopamine has been on a downward trend(**E**). DEPC: *Aedes aegypti* injected with DEPC water; *ds GUS*: *Aedes aegypti* injected with the GUS dsRNA; *ds aaNAT1* 2h: *Aedes aegypti* two hours after *AaaaNAT1* ds RNA injection; *ds aaNAT1* 6h: *Aedes aegypti* six hours after *AaaaNAT1* ds RNA injection; *ds aaNAT1* 12h: *Aedes aegypti* twelve hours after *AaaaNAT1* ds RNA injection. \*\*\*\**p* <0.0001, *** *p* <0.001, ** *p* <0.01, * *p* <0.05.

**S1 Table. List of primers for qPCR.**

## Notes

### Competing Interest Statement

The authors have declared no competing interest.

## References

1. Guo C, Zhou Z, Wen Z, Liu Y, Zeng C, Xiao D, et al. Global Epidemiology of Dengue Outbreaks in 1990–2015: A Systematic Review and Meta-Analysis. Front Cell Infect Microbiol. 2017;7: 317. doi:10.3389/fcimb.2017.00317

2. Ribeiro JMC, Arc à B. Chapter 2 From Sialomes to the Sialoverse. Advances in Insect Physiology. Elsevier; 2009. pp. 59–118. doi:10.1016/S0065-2806(09)37002-2

3. Ribeiro JMC. Role of Saliva in Blood-Feeding by Arthropods. Annu Rev Entomol. 1987;32: 463–478. doi:10.1146/annurev.en.32.010187.002335

4. Mehere P, Han Q, Christensen BM, Li J. Identification and characterization of two arylalkylamine N-acetyltransferases in the yellow fever mosquito, Aedes aegypti. Insect Biochemistry and Molecular Biology. 2011;41: 707–714. doi:10.1016/j.ibmb.2011.05.002

5. Klein DC. Evolution of The Vertebrate Pineal Gland: The Aanat Hypothesis. Chronobiology International. 2006;23: 5–20. doi:10.1080/07420520500545839

6. Klein DC. Arylalkylamine N-Acetyltransferase: “the Timezyme. ” Journal of Biological Chemistry. 2007;282: 4233–4237. doi:10.1074/jbc.R600036200

7. Davla S, Artiushin G, Li Y, Chitsaz D, Li S, Sehgal A, et al. AANAT1 functions in astrocytes to regulate sleep homeostasis. Elife. 2020;9: e53994. doi:10.7554/eLife.53994

8. Hiragaki S, Suzuki T, Mohamed AAM, Takeda M. Structures and functions of insect arylalkylamine N-acetyltransferase (iaaNAT); a key enzyme for physiological and behavioral switch in arthropods. Front Physiol. 2015;6. doi:10.3389/fphys.2015.00113

9. Noh MY, Koo B, Kramer KJ, Muthukrishnan S, Arakane Y. Arylalkylamine N-acetyltransferase 1 gene (TcAANAT1) is required for cuticle morphology and pigmentation of the adult red flour beetle, Tribolium castaneum. Insect Biochemistry and Molecular Biology. 2016;79: 119–129. doi:10.1016/j.ibmb.2016.10.013

10. Wang Z-X, Liu Y-L, Teng F-Y, Lu Y-Y, Qi Y-X. Arylalkylamine N-acetyltransferase 1 gene (AANAT1) regulates cuticle pigmentation and ovary development of the adult oriental fruit fly, Bactrocera dorsalis. Insect Biochem Mol Biol. 2022;150: 103850. doi:10.1016/j.ibmb.2022.103850

11. Richardt A, Kemme T, Wagner S, Schwarzer D, Marahiel MA, Hovemann BT. Ebony, a Novel Nonribosomal Peptide Synthetase for β-Alanine Conjugation with Biogenic Amines in Drosophila. Journal of Biological Chemistry. 2003;278: 41160–41166. doi:10.1074/jbc.M304303200

12. Liao C, Upadhyay A, Liang J, Han Q, Li J. 3,4-Dihydroxyphenylacetaldehyde synthase and cuticle formation in insects. Developmental & Comparative Immunology. 2018;83: 44–50. doi:10.1016/j.dci.2017.11.007

13. Han Q, Robinson H, Ding H, Christensen BM, Li J. Evolution of insect arylalkylamine *N* - acetyltransferases: Structural evidence from the yellow fever mosquito, *Aedes aegypti*. Proc Natl Acad Sci USA. 2012;109: 11669–11674. doi:10.1073/pnas.1206828109

14. Dempsey DR, Jeffries KA, Bond JD, Carpenter A-M, Rodriguez-Ospina S, Breydo L, et al. Mechanistic and Structural Analysis of *Drosophila melanogaster* Arylalkylamine *N* - Acetyltransferases. Biochemistry. 2014;53: 7777–7793. doi:10.1021/bi5006078

15. Barberà M, Mengual B, Collantes-Alegre JM, Cortés T, González A, Martínez-Torres D. Identification, characterization and analysis of expression of genes encoding arylalkylamine N -acetyltransferases in the pea aphid *A cyrthosiphon pisum*. Insect Molecular Biology. 2013;22: 623–634. doi:10.1111/imb.12050

16. Zhang L, Tang Y, Chen H, Zhu X, Gong X, Wang S, et al. Arylalkalamine *N* - acetyltransferase-1 acts on a secondary amine in the yellow fever mosquito, *Aedes aegypti*. FEBS Letters. 2022;596: 1081–1091. doi:10.1002/1873-3468.14316

17. Hiragaki S, Suzuki T, Mohamed AAM, Takeda M. Structures and functions of insect arylalkylamine N-acetyltransferase (iaaNAT); a key enzyme for physiological and behavioral switch in arthropods. Front Physiol. 2015;6. doi:10.3389/fphys.2015.00113

18. Zeng T, Teng F, Wei H, Lu Y, Xu Y, Qi Y. AANAT1 regulates insect midgut detoxification through the ROS/CncC pathway. Commun Biol. 2024;7: 808. doi:10.1038/s42003-024-06505-x

19. Braverman IM. The Cutaneous Microcirculation: Ultrastructure and Microanatomical Organization. Microcirculation. 1997;4: 329–340. doi:10.3109/10739689709146797

20. Hundelshausen P, Petersen F, Brandt E. Platelet-derived chemokines in vascular biology. Thromb Haemost. 2007;97: 704–713. doi:10.1160/TH07-01-0066

21. Nuttall GA. Hemostasis and Thrombosis: Basic Principles and Clinical Practice, 5th ed. Anesthesia & Analgesia. 2007;104: 1317. doi:10.1213/01.ANE.0000263681.99710.0B

22. Nemerson Y. Tissue factor and hemostasis. Blood. 1988;71: 1–8.

23. Taracena M, Hunt C, Pennington P, Andrew D, Jacobs-Lorena M, Dotson E, et al. Effective Oral RNA Interference (RNAi) Administration to Adult Anopheles gambiae Mosquitoes. JoVE. 2022; 63266. doi:10.3791/63266

24. Suski J, Lebiedzinska M, Bonora M, Pinton P, Duszynski J, Wieckowski MR. Relation Between Mitochondrial Membrane Potential and ROS Formation. In: Palmeira CM, Moreno AJ, editors. Mitochondrial Bioenergetics. New York, NY: Springer New York; 2018. pp. 357–381. doi:10.1007/978-1-4939-7831-1_22

25. Nolfi-Donegan D, Braganza A, Shiva S. Mitochondrial electron transport chain: Oxidative phosphorylation, oxidant production, and methods of measurement. Redox Biology. 2020;37: 101674. doi:10.1016/j.redox.2020.101674

26. Calvo E, Mizurini DM, Sá-Nunes A, Ribeiro JMC, Andersen JF, Mans BJ, et al. Alboserpin, a Factor Xa Inhibitor from the Mosquito Vector of Yellow Fever, Binds Heparin and Membrane Phospholipids and Exhibits Antithrombotic Activity. Journal of Biological Chemistry. 2011;286: 27998–28010. doi:10.1074/jbc.M111.247924

27. Tschuor C, Asmis LM, Lenzlinger PM, Tanner M, Härter L, Keel M, et al. In vitro norepinephrine significantly activates isolated platelets from healthy volunteers and critically ill patients following severe traumatic brain injury. Crit Care. 2008;12: R80. doi:10.1186/cc6931

28. Belyaev AV, Dunster JL, Gibbins JM, Panteleev MA, Volpert V. Modeling thrombosis in silico: Frontiers, challenges, unresolved problems and milestones. Physics of Life Reviews. 2018;26–27: 57–95. doi:10.1016/j.plrev.2018.02.005

29. Bye AP, Unsworth AJ, Gibbins JM. Platelet signaling: a complex interplay between inhibitory and activatory networks. Journal of Thrombosis and Haemostasis. 2016;14: 918–930. doi:10.1111/jth.13302

30. Farjana T, Tuno N. Multiple blood feeding and host-seeking behavior in Aedes aegypti and Aedes albopictus (Diptera: Culicidae). J Med Entomol. 2013;50: 838–846. doi:10.1603/me12146

31. Georgiades M, Alampounti A, Somers J, Su MP, Ellis DA, Bagi J, et al. Hearing of malaria mosquitoes is modulated by a beta-adrenergic-like octopamine receptor which serves as insecticide target. Nat Commun. 2023;14: 4338. doi:10.1038/s41467-023-40029-y

32. Lapshin DN, Vorontsov DD. Functions of the Auditory System of Female Mosquitoes (Diptera, Culicidae). Entmol Rev. 2023;103: 251–262. doi:10.1134/S0013873823030016

33. Finetti L, Paluzzi J-P, Orchard I, Lange AB. Octopamine and tyramine signalling in Aedes aegypti: Molecular characterization and insight into potential physiological roles. Hull JJ, editor. PLoS ONE. 2023;18: e0281917. doi:10.1371/journal.pone.0281917

34. Klein DC, Coon SL, Roseboom PH, Weller JL, Bernard M, Gastel JA, et al. The melatonin rhythm-generating enzyme: molecular regulation of serotonin N-acetyltransferase in the pineal gland. Recent Prog Horm Res. 1997;52: 307–357; discussion 357-358.

35. Zilberman-Peled B, Ron B, Gross A, Finberg JPM, Gothilf Y. A possible new role for fish retinal serotonin-N-acetyltransferase-1 (AANAT1): Dopamine metabolism. Brain Research. 2006;1073–1074: 220–228. doi:10.1016/j.brainres.2005.12.028

36. Iuvone PM, Chong NW, Bernard M, Brown AD, Thomas KB, Klein DC. Melatonin biosynthesis in chicken retina. Regulation of tryptophan hydroxylase and arylalkylamine N-acetyltransferase. Adv Exp Med Biol. 1999;460: 31–41.

37. Falcón J, Bolliet V, Collin JP. Partial characterization of serotonin N - acetyltransferases from northern pike (Esox lucius, L.) pineal organ and retina: effects of temperature. Pflugers Arch. 1996;432: 386–393. doi:10.1007/s004240050149

38. Mizusawa K, Iigo M, Suetake H, Yoshiura Y, Gen K, Kikuchi K, et al. Molecular Cloning and Characterization of a cDNA Encoding the Retinal Arylalkylamine N-Acetyltransferase of the Rainbow Trout, Oncorhynchus mykiss. Zoolog Sci. 1998;15: 345–351. doi:10.2108/zsj.15.345

39. Bégay V, Falcón J, Cahill GM, Klein DC, Coon SL. Transcripts encoding two melatonin synthesis enzymes in the teleost pineal organ: circadian regulation in pike and zebrafish, but not in trout. Endocrinology. 1998;139: 905–912. doi:10.1210/endo.139.3.5790

40. Gothilf Y, Coon SL, Toyama R, Chitnis A, Namboodiri MA, Klein DC. Zebrafish serotonin N-acetyltransferase-2: marker for development of pineal photoreceptors and circadian clock function. Endocrinology. 1999;140: 4895–4903. doi:10.1210/endo.140.10.6975

41. Puppala D, Maaswinkel H, Mason B, Legan SJ, Li L. An in vivo microdialysis study of light/dark-modulation of vitreal dopamine release in zebrafish. J Neurocytol. 2004;33: 193–201. doi:10.1023/b:neur.0000030694.88653.d6

42. Barberà M, Mengual B, Collantes-Alegre JM, Cortés T, González A, Martínez-Torres D. Identification, characterization and analysis of expression of genes encoding arylalkylamine N-acetyltransferases in the pea aphid Acyrthosiphon pisum. Insect Mol Biol. 2013;22: 623– 634. doi:10.1111/imb.12050

43. Bembenek J, Sehadova H, Ichihara N, Takeda M. Day/night fluctuations in melatonin content, arylalkylamine N-acetyltransferase activity and NAT mRNA expression in the CNS, peripheral tissues and hemolymph of the cockroach, Periplaneta americana. Comp Biochem Physiol B Biochem Mol Biol. 2005;140: 27–36. doi:10.1016/j.cbpc.2004.03.017

44. Hardie J, Gao N. Melatonin and the pea aphid, Acyrthosiphon pisum. J Insect Physiol. 1997;43: 615–620. doi:10.1016/s0022-1910(97)00015-2

45. Hintermann E, Grieder NC, Amherd R, Brodbeck D, Meyer UA. Cloning of an arylalkylamine N-acetyltransferase (aaNAT1) from Drosophila melanogaster expressed in the nervous system and the gut. Proc Natl Acad Sci U S A. 1996;93: 12315–12320. doi:10.1073/pnas.93.22.12315

46. Itoh MT, Hattori A, Nomura T, Sumi Y, Suzuki T. Melatonin and arylalkylamine N-acetyltransferase activity in the silkworm, Bombyx mori. Mol Cell Endocrinol. 1995;115: 59–64. doi:10.1016/0303-7207(95)03670-3

47. Itoh MT, Hattori A, Sumi Y, Suzuki T. Day-night changes in melatonin levels in different organs of the cricket (Gryllus bimaculatus). J Pineal Res. 1995;18: 165–169. doi:10.1111/j.1600-079x.1995.tb00156.x

48. Choo Y-M, Buss GK, Tan K, Leal WS. Multitasking roles of mosquito labrum in oviposition and blood feeding. Front Physiol. 2015;6: 306. doi:10.3389/fphys.2015.00306

49. Kraemer MUG, Sinka ME, Duda KA, Mylne A, Shearer FM, Brady OJ, et al. The global compendium of Aedes aegypti and Ae. albopictus occurrence. Sci Data. 2015;2: 150035. doi:10.1038/sdata.2015.35

50. Abramson DI, Allen JC, Baden HP, Benyon RC, Bird AG, Brain SD, et al. Pharmacology of Skin Systems Autocoids in Normal and Inflamed Skin.

51. Tagami T. Antithrombin concentrate use in sepsis-associated disseminated intravascular coagulation: re-evaluation of a “pendulum effect” drug using a nationwide database. J Thromb Haemost. 2018;16: 458–461. doi:10.1111/jth.13948

52. Chen J, Lu H-R, Zhang L, Liao C-H, Han Q. RNA interference-mediated knockdown of 3, 4-dihydroxyphenylacetaldehyde synthase affects larval development and adult survival in the mosquito Aedes aegypti. Parasites Vectors. 2019;12: 311. doi:10.1186/s13071-019-3568-7

53. Taracena M, Hunt C, Pennington P, Andrew D, Jacobs-Lorena M, Dotson E, et al. Effective Oral RNA Interference (RNAi) Administration to Adult Anopheles gambiae Mosquitoes. JoVE. 2022; 63266. doi:10.3791/63266

